# Polygenic adaptation to overnutrition reveals a role for cholinergic signaling in longevity

**DOI:** 10.1101/2023.06.14.544888

**Authors:** Thomas B Rundell, Melina Brunelli, Azva Alvi, Gabrielle Safian, Christina Capobianco, Wangshu Tu, Sanjeena Subedi, Anthony Fiumera, Laura Palanker Musselman

## Abstract

Overnutrition by high-sugar (HS) feeding reduces both the lifespan and healthspan across taxa. Pressuring organisms to adapt to overnutrition can highlight genes and pathways important for the healthspan in stressful environments. We used an experimental evolution approach to adapt four replicate, outbred population pairs of *Drosophila melanogaster* to a HS or control diet. Sexes were separated and aged on either diet until mid-life, then mated to produce the next generation, allowing enrichment for protective alleles over time. All HS-selected populations increased their lifespan and were therefore used as a platform to compare allele frequencies and gene expression. Pathways functioning in the nervous system were overrepresented in the genomic data and showed evidence for parallel evolution, although very few genes were the same across replicates. Acetylcholine-related genes, including the muscarinic receptor *mAChR-A,* showed significant changes in allele frequency in multiple selected populations and differential expression on a HS diet. Using genetic and pharmacological approaches, we show that cholinergic signaling affects Drosophila feeding in a sugar-specific fashion. Together, these results suggest that adaptation produces changes in allele frequencies that benefit animals under conditions of overnutrition and that it is repeatable at the pathway level.

## Introduction

The adaptation of complex phenotypes arises from a composite of physiological and morphological traits under simultaneous selection (1,2). Aging is one complex trait of interest that is accompanied by pathophysiology across a wide variety of outcomes (3). In many cases, long-lived individuals also display increases in the healthspan, and age is a risk factor for a range of pathophysiologies (4–7). Although it is generally true that the longest-lived in a population display healthier quantitative traits than short-lived, the correlation between traits can be weak and the mechanisms that underlie each correlation are not well understood (8,9). In addition, such traits as aging are highly polygenic, with many loci of small effect sizes contributing to the phenotype. The polygenic nature of complex traits creates challenges for researchers attempting to map the genetic basis of said traits (1,2,10–13). One approach to address this question that has met with success has been evolve-and-resequence (E&R) studies using model organisms such as *Drosophila melanogaster* (abbreviated as *Drosophila* here). These studies have used a variety of selection pressures on life-history traits such as increased longevity (11,14) and late-life reproduction (15–21) to identify loci that increase lifespan. An advantage of E&R studies in uncovering physiologically relevant adaptation is the ability to leverage replicate populations from a common genetic background. Parallelism across replicates thus increases confidence in potentially causal mechanisms at the SNP, gene, and/or pathway levels.

A poor diet, or dietary excess, often accelerates the pathophysiology phenotypes associated with senescence. Overnutrition contributes to so-called metabolic diseases such as type 2 diabetes (T2D) and cardiovascular disease, which are thought to be caused by the interaction of genetic susceptibility with the environment (22). It has been well established that sedentary lifestyles and diets high in sugar and fat are primary risk factors in the development of metabolic disease (22–24). The analogous pathophysiology has been successfully modeled in several animal models, including mice and *Drosophila*, using high-sugar or other high-calorie diets (25–33). Flies fed a high-sugar (HS) diet exhibit a reduced lifespan and healthspan, including obesity, hyperglycemia, and impaired immunity, reproduction, feeding behavior, insulin signaling, and cardiac function (26,28,30,32,34–37). As complex traits, the genetic susceptibilities to metabolic disease are thought to be highly polygenic with SNPs of low effect sizes (22). GWA studies have previously been used to identify such SNPs that have been validated in model organisms, including *Drosophila* (38–40). Despite the success of such approaches at identifying linked loci, the effect sizes of these SNPs are insufficient to fully capture the genetic component of disease susceptibility (23).

In the present study, we have leveraged the negative health outcomes of HS-induced overnutrition as a selective pressure to enrich the fly genome for protective alleles via experimental evolution. We show that genetically diverse populations of *Drosophila* have significantly improved lifespans on high-sugar and control diets after 10 generations of selection. These increases correlated with significant changes in allele frequencies and expression profiles of a number of genes, many of which are related to cholinergic signaling. We found that a reduction in one gene with significant alterations in allele frequencies and expression, the muscarinic acetylcholine receptor type-A (*mAChR-A*), reduces lifespan and feeding rates in a sugar-specific manner.

## Methods

### Husbandry for experimental evolution

Large populations were reared and aged with controlled humidity at 25°C on a 12:12 light:dark cycle. Both 0.15M (5%) sucrose control media and 1M (34%) sucrose selective media received 3x tegosept mold inhibitor but was otherwise as described previously (41). Both control and selection populations were reared on 0.15M media and then aged in (30cm^3) BugDorm insect rearing cages (MegaView Science Co., Ltd., Taiwan Cat. #DP1000). Control populations were aged on 0.15M media while selection populations were aged on the selective 1M sucrose diet.

### High-sugar selection regime

The starter population was obtained as a tray of 16 bottles from the Julien Ayroles lab at Princeton University, where 40 wild-caught Netherlands isofemale inbred lines were crossed in round robin fashion to produce a large, outbred population as previously described (42). This starter population was then divided into 8 populations. These 8 populations were then randomly paired such that each pair of Control (C1-4) and Selection (S1-4) populations shared the same incubator space and developmental timing. For every generation, the HS population was allowed to mate for 3 days after eclosion before being separated by sex into cages. Census populations were generated at a size of ∼6000 flies, 3000 males and 3000 females. Both sexes were aged at 25°C in 12:12 light:dark conditions on HS food until ∼50% of the population died off (paired control typically exhibited 10-20% mortality at this time point), after which the male and female cages were joined to allow mating. Eggs were laid on LS food and bottles were removed from the cages for development. Upon eclosion, adults were allowed to mate for 3 days before transfer to the Control or Selective diets. These adults then constituted the next non-overlapping generation. As Drosophila display nearly complete last male sperm precedence after significant passages of time, we do not expect that the 3-day mating affected offspring paternity (43,44). Offspring under3-daydevelopment on control food (0.15 M sucrose) to avoid the detrimental developmental effects of rearing larvae in HS media (35,36). HS selection was thus confined to adult feeding. Control populations mimicked the timing of the HS population with which they were paired, such that when the paired HS population reached 50% death and underwent mating, so too did the control population (typically at 10-20% death).

### Fly lines and husbandry for genetics

TRiP insertion site control genotype (BDSC stock #36303) and the *UAS-mAChR-a* RNAi (BDSC stock #27571) stock lines were obtained from the Bloomington Drosophila Stock Center (NIH P40OD018537). Flies were reared and aged with controlled humidity at 25°C on a 12:12 light:dark cycle. Fly stocks were reared on a 5% dextrose-cornmeal-yeast-agar media diet. The *mAChR-a* RNAi line was crossed with *UAS-Dcr2;*syb-GAL4 (45) to facilitate the brain-specific knockdown of *mAChR-a*. Crosses were reared on 5% dextrose-cornmeal-yeast-agar media and within 24 hrs of eclosion transferred onto either 1M sucrose or 0.15M sucrose and aged for 1 or 3 weeks.

### Atropine administration

The muscarinic receptor antagonist atropine ((*RS*)-(8-Methyl-8-azabicyclo[3.2.1]oct-3-yl, VWR Cat. TCA0754-005G) was dissolved in ethanol and added to 1M (HS) or 0.15M (LS) sucrose food media at a final concentration of 0.1% atropine as previously described (46,47). The control media had ethanol added to the same final concentration without atropine. Mated flies aged 1-3 days were flipped onto atropine containing food and aged for 3 weeks.

### Survival/aging

Mated male and female flies were collected 3 days after eclosion and aged separately on control and experimental diets at a density of 30 flies/vial. Food media was provided with watered chromatography paper wicks to ensure flies had sufficient water. Food was replaced and surviving flies were counted every 2-3 days. Each group consisted of 120 flies. Kaplan-Meier curves were generated, and curves were compared using a Mantel-Cox logrank test. Error bars represent standard error calculated using the Greenwood method. Median day of death was calculated as the time at which survival probability was equal to 50%. Computations were conducted in GraphPad Prism software (v. 9.4.0).

### DNA isolation and sequencing

At generations 0, 5, 10, and 15, pools of 120 flies per replicate were collected and frozen at - 80°C until processing (48,49). Frozen flies were homogenized in PBS (pH 7.4) and DNA was extracted per the manufacturer’s instructions using Qiagen DNeasy Blood and Tissue kit (Qiagen, Germantown, MD, USA Cat. #69504). Samples were then subjected to ethanol precipitation as previously described (50). Total DNA was quantified using Qubit 2.0 fluorometer HS Assay kit (Life Technologies/ThermoFisher Scientific, Waltham, MA, USA Cat. #32850). Libraries were constructed and sequencing was performed by the Genome Technology Access Center at the McDonnell Genome Institute (Washington University in St. Louis, Missouri). DNA was sequenced on the Illumina NovaSeq 6000 platform and mapped to BDGP release 6.27. Generation 0 had an average number of mapped reads of over 332,000,000 at an average 88% mapped reads. Generation 0 had an average read depth of >320x. Generation 10 had an average number of mapped reads over 320,000,000 and an average 92% mapped reads to give a sequencing depth of >240x across all populations.

### RNA isolation and sequencing

At generation 10, flies were aged on 1M sucrose food media for three weeks and frozen in groups of 20 per replicate at −80°C until processing. Flies were then homogenized by electric mortar and pestle in Ribozol (VWR Cat. N580-100ML). RNA was treated with DNase (VWR Cat. PIER89836) and then extracted per the manufacturer’s instructions using the Qiagen RNeasy Mini Kit (Qiagen, Germantown, MD, USA Cat #74104). Total RNA was quantified using Qubit 2.0 fluorometer HS Assay kit (Life Technologies/ThermoFisher Scientific, Waltham, MA, USA Cat. # 10210). Libraries were constructed after FastSelect (Qiagen, Germantown, MD, USA Cat. #334385) rRNA subtraction and sequencing was performed by the Genome Technology Access Center at the McDonnell Genome Institute (St. Louis, Missouri) using an Illumina HiSeq platform. All samples produced over 11,000,000 mapped reads with an average of 12,360,000 mapped reads and an average 93% mapped reads. RNA reads were aligned to the Ensembl release 76 genome using STAR (v. 2.5.1a) (51).

### Gene identification

To identify SNPs that were likely to have changed in allele frequency as a result of selection to high-sugar, we used Popoolation2 (48). Popoolation2’s Fisher’s exact test was used to identified SNPs significantly differentiated in allele frequency between selected populations at generation 10 and the starter population at generation 0 with a Bonferroni corrected P value of 0.05 (raw P vulaues of 0.05 adjusted to ∼10^-8^. To eliminate the effects of selection to other sources (e.g. lab rearing conditions) we further identified SNPs that differentiated between the selected populations at generation 10 and the control populations at generation 10 with a Bonferroni corrected P value of 0.05. Genes of interest were called from SNPs if at least one significant SNP was within an annotated gene or within 2 kb upstream or downstream of a gene. In combination with the DNA analysis after selection, we conducted a differential expression analysis on generation 10 RNA-seq data. Gene counts were normalized using the TMM method in the R package EdgeR (52). Differential expression analysis was then conducted using the R package Limma and Benjamini-Hochberg procedure was used for multiple testing correction (53). Comparisons were between Selected vs Control population pairs at generation 10. Genes with false-discovery rate adjusted P values less than or equal to 0.05 were considered differentially expressed. Version 4.0.5 was used for all analyses in R.

### Cluster analysis

Model-based clustering was utilized for cluster analysis. Model-based clustering assumes that the population comprises of subpopulations or clusters and typically each cluster is represented by a probability density. Here, we used a mixture of multivariate Poisson log-normal distributions in which each cluster is modelled using a multivariate Poisson log-normal distribution, a distribution ideal for modelling multivariate count data arising from RNAseq studies (54). Several models with varying number of components/clusters were fitted to the dataset and the best fitting model was selected using a model-selection criteria.

### Network analysis

To understand more about the target gene mAChR-A and its function, we performed a network analysis to learn its relationship with other genes that have been previously studied. First, we conducted a gene ontology (GO) analysis using gProfiler to select the genes that were significant and associated with GO terms of interest, such as DNA binding, and ribosome biogenesis (55). 360 genes were selected from the male group. We then used GeneMANIA (56) to identify 8 genes that were directly linked to mAChR-A and utilized Cytoscape (57) to produce a network plot which contains an overall network and the types of interactions among these 9 genes.

### Feeding assay

Feeding assays were performed similarly to those described in (58). Flies were aged for 1 or 3 weeks on control and experimental food media then transferred to corresponding media supplemented with 2% FD&C Blue #1 for 2 hrs (11am-1pm) and frozen immediately at −80°C until processing. Flies were homogenized in groups of 4 with mortar and pestle and supernatants were read at 630 nm on a VersaMax microplate spectrophotometer (Molecular Devices). Statistical analyses were conducted using a two-tailed Student’s *t*-test or one-way ANOVA on Prism software (v. 9.4.0).

## Results

### Experimental selection to a high-sugar-diet increased longevity on control and selective diets

To identify genes that increase lifespan on a high-sugar (HS) diet, we undertook an experimental evolution approach. Large census populations (∼6000 flies) were reared on control (LS) diets, then aged on LS or HS in single-sex cages until approximately half the flies in the selected population had died (Fig 1; see methods for more detail). The starting population was a diverse outbred population created by round-robin mating of wild-caught isofemale lines (citation; see methods for more details). Replicate populations were paired as shown (Fig 1) and matched in husbandry and developmental timing.

**Fig 1.**
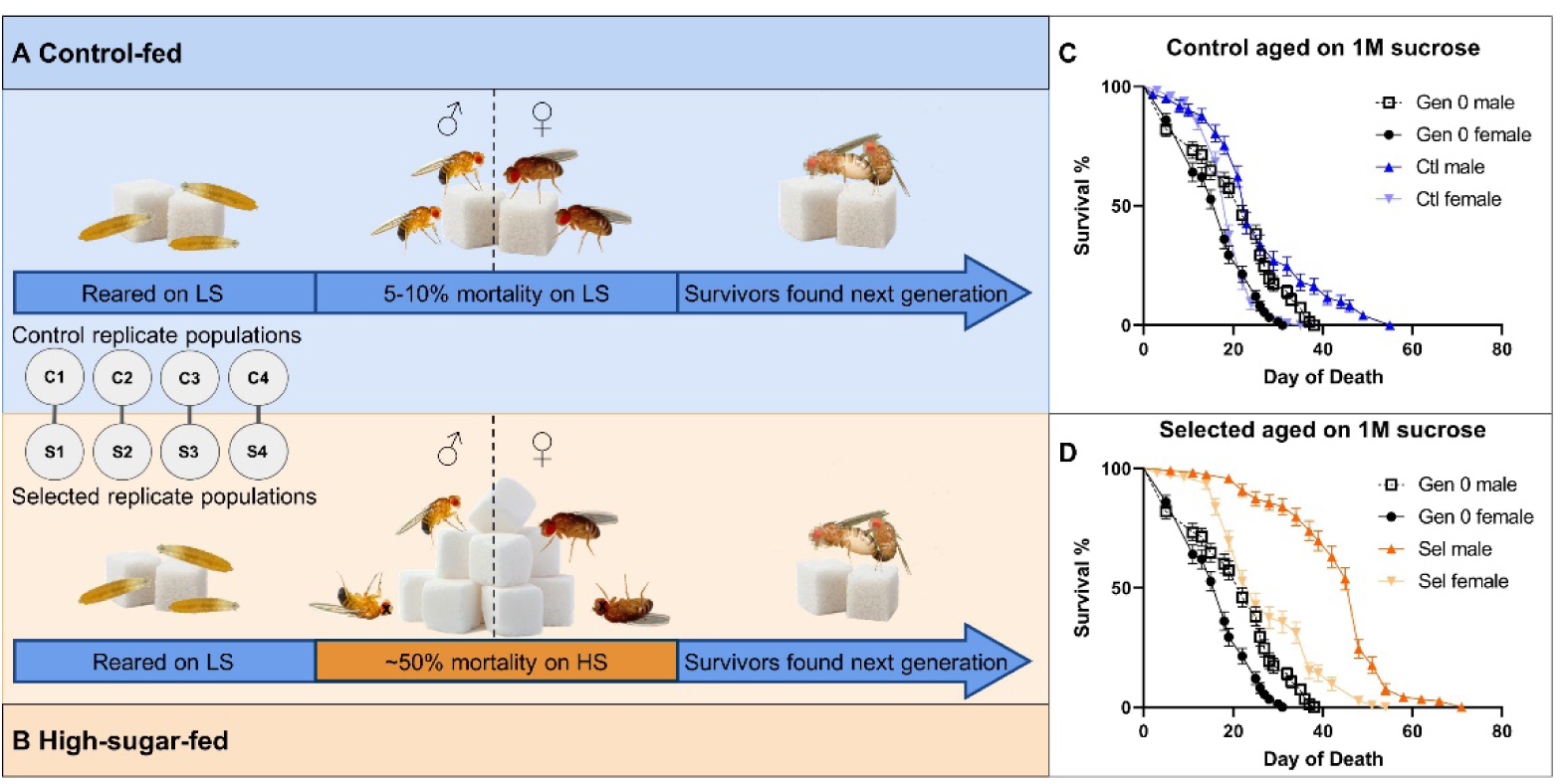
Schematic for adaptation to high-sugar feeding. Eight replicate populations with a census population of ∼6000 flies were paired such that one LS and HS pair experienced the same developmental timing and laboratory conditions. Mated males and females were aged separately on (A) 0.15M sucrose for LS populations and (B) 1M sucrose for HS populations. HS flies were aged until ∼50% mortality. The survivors were then used to found the subsequent generation. In parallel, the LS populations were each mated at the same time as their paired HS populations. (C) Most control-adapted populations increased in lifespan, but this benefit was small in magnitude on the HS diet. (D) HS-selected populations had larger increases in lifespan on the HS diet.

In response to 10 generations of selection to adult feeding on a high-sugar diet, all Selected populations exhibited a significant increase in longevity relative to both generation 0 (G0) and to the paired control population when challenged with the selective HS (Fig. 2A-B,E-F). Interestingly, selection to HS adult feeding also extended lifespan in most populations on the non-selective, LS diet. It is also notable that all control populations showed an increase in lifespan on LS compared to generation 0, suggesting that even at a modest 10-15% mortality rate selection for increased lifespan occurred (Fig 2A-B, E-F).

**Fig 2.**
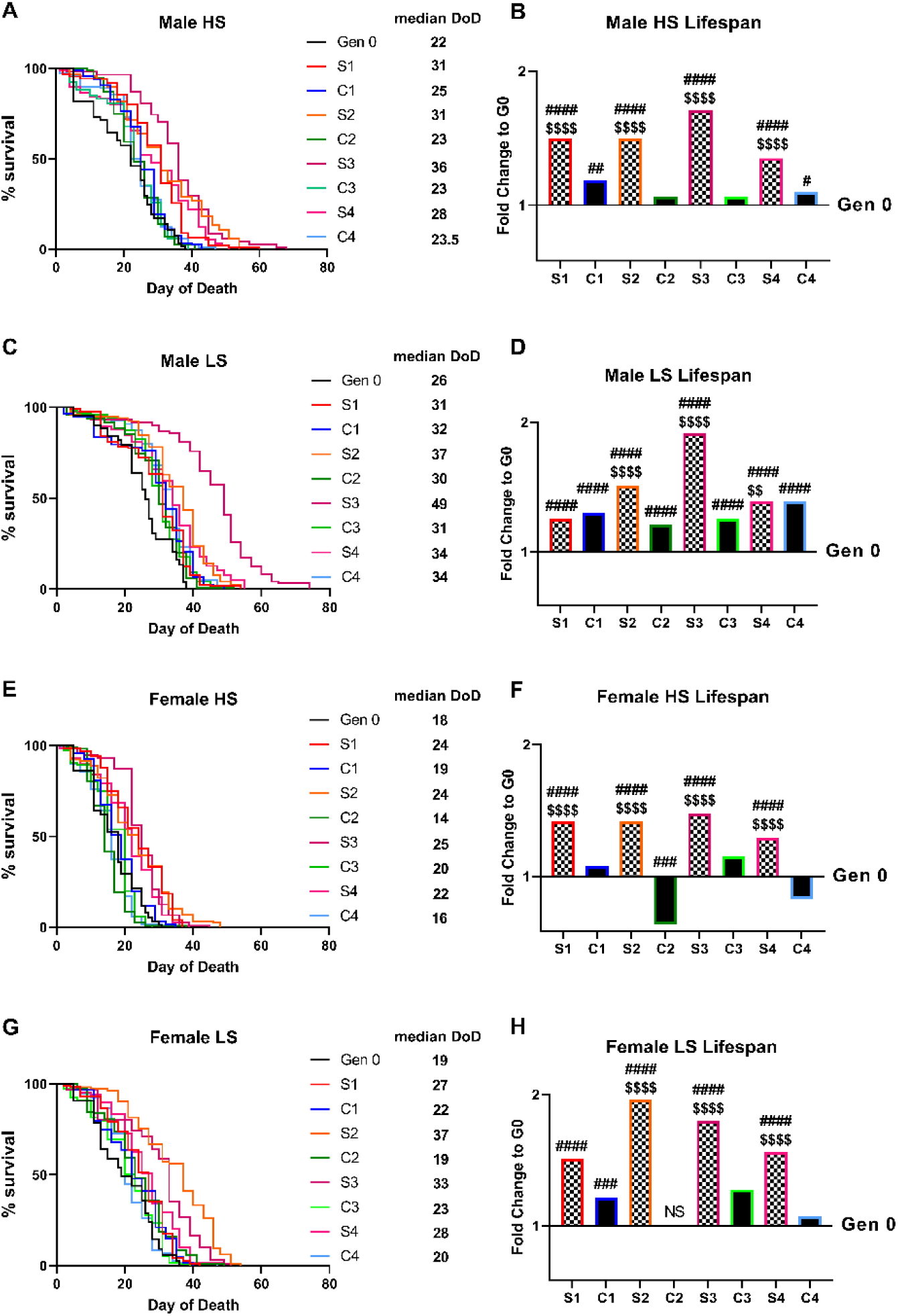
Increased longevity in all populations after selection. In cohorts of 30, 120 recently eclosed mated males (A-D) or females (E-H) were aged on 0.15M (LS) or 1M (HS) sucrose diets. (A, C, E, G) Survival curves for control populations are shown in cool colors, high-sugar selected populations in warm colors. Median day of death (DoD) is shown for each population alongside the survival curve legend. (B, D, F, H) Bars represent log_2_ fold change of median day of death compared to generation 0 median day of death on respective diets. The significance corresponding to number of symbols is as follows: *P < 0.5, **P < 0.01, ***P < 0.001, ****P < 0.0001 by log-rank Mantel-Cox. # represents the significance for generation 10 vs generation 0. $ represents the significance for Selected population vs Control population at generation 10.

Interestingly, the magnitude of changes in lifespan were sex-specific, with Selected males exhibiting greater increases in survival on HS (1.74-fold increase in median day of death compared to the control 1.1-fold increase). By contrast, selected females exhibited an average fold-increase of 1.34 compared to the control populations’ 1.01. It is notable that males exhibited increased lifespan compared to females in all populations on both diets.

Replicate populations, despite having a common starting genotype, did not respond to selection in the same was at the genotype or phenotype level. While all selected populations had increased lifespan relative to their paired control on HS diets, not all exhibited such increases on LS feeding. Notably, S2’s lifespan did not significantly differ from its paired control in males or females on LS. Populations S3 and S4, by contrast, stand out for their drastic increase in lifespan across all diets and in both sexes. Indeed, these two populations both exhibit the highest fold changes on HS food and exceed a 2-fold increase in median day of death compared to generation 0 on LS diets in both sexes (Figure 2, S1-4 Fig). Population C4 also stands out, as it shows no increase in survival on HS in females, but a modest increase on HS in males.

### Significant effects on allele frequencies after 10 generations of selection

To identify genes of interest that may have conferred an adaptive benefit during HS feeding in the selected populations, we utilized next-generation pool-sequencing (Pool-seq) (48) at generation 0 and after generation 10 to estimate population-wide allele frequencies in pools of 120 males and females (60 males, 60 females). Pool-seq allows us to obtain deep sequencing (∼average 320x coverage) to identify population-wide changes that correlate with increased lifespan in response to our HS selective pressure (48). Candidate SNPs were then identified using a pairwise Fisher’s exact test with a Bonferroni corrected threshold for multiple tests of *P* < 0.05 (Fig3; S5 Fig). Using these criteria, we identified between 89,909 and 121,850 different SNPs from generation 0 to generation 10 in the HS-adapted S1, S2, S3, and S4 populations. There were fewer SNPs that changed significantly in the control populations, from 41,934 to 104,976 different SNPs, in the control populations C1, C2, C3, and C4. Significant loci showed patterns of clustering across the genome, suggestive of both polygenic adaptation and the presence of hitchhiking loci linked to local, causal SNPs (S5 Fig). We next compared the control and selected population pairs directly using a pairwise Fisher’s exact test with a Bonferroni corrected threshold for multiple tests of P < 0.05. Here we identified 32,100 SNPs that differed between the S1-C1 pairing, 5,574 between S2-C2, 10,111 between S3-C3, and 20,001 between the S4-C4 pairing. SNPs were then further filtered to those within a gene or within 2 kb upstream or downstream of a gene. While there was a sizable proportion of genes with overlap between at least two selected populations (21%), there still existed a sizable proportion of genes unique to each evolved population (79%) (Fig 3A). Interestingly, there was a much stronger overlap in significant gene ontological (GO) categories as compared to the individual genes (Fig 3B,C). This suggests strong evidence for parallel evolution in the pathways that are involved but that unique genes within those pathways responded to selection in the different replicate populations. One such class of neuronal categories that were consistent across populations included learning/memory, generation of neurons, neuronal development, GPCR signaling, and behavior (-log_10_(P) > 7.6 for all populations, Fig 3C, S6 Fig). The top GO tag in each population pair comparison was involved in neural development (S6 Fig).

**Fig 3.**
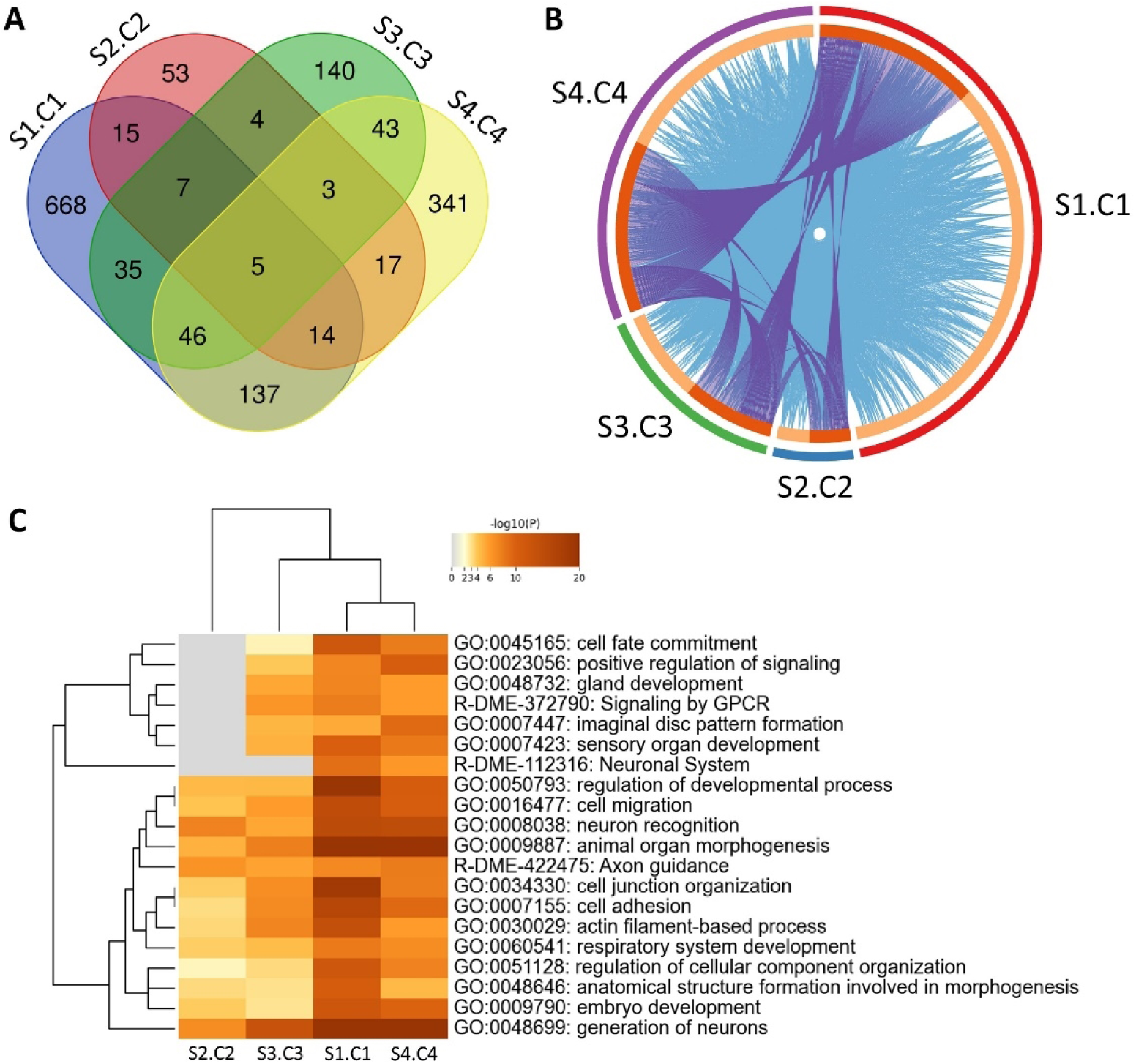
Pooled whole-genome sequencing shows changes in allele frequencies after ten generations of selection. (A) Venn diagram showing overlap in number of genes containing (or within 2 kb upstream or downstream of the gene) at least one significant SNP by Fisher’s exact test between Selected and Control population pairs. 21% of identified genes overlap between at least two populations, with a majority of identified genes being unique to a selected population pair (79%). The 5 genes identified in all Selected populations were *rhea* (FBgn0260442)*, cdi* (FBgn0004876)*, kst* (FBgn0004167)*, SKIP* (FBgn0051163), and *CG7720* (FBgn0038652). (B) Circos plot showing overlap of significant identical genes (purple lines) and genes with the same ontology tag (blue lines). Terms with P < 0.01 and enrichment score >1.5 were counted as significant categories (C) Heatmap with hierarchical clustering of significant ontological terms identified in each respective population pair. Heatmap coloring depicts significant P values. Terms with P < 0.01 and enrichment score >1.5 were counted as significant categories.

### Dynamic gene expression profiles observed across selected population pairs

We next measured changes in gene expression between selected and control populations after 10 generations of selection. Because expression profiles differed so much between males and females, we analyzed these datasets separately (Fig 4A-B, S7 Fig). Differentially expressed genes were identified in a pair-wise fashion, comparing each Selected population to its Control population partner. Genes with false-discovery rate adjusted P values less than or equal to 0.05 were considered differentially expressed. Transcriptomes differed between Selected and Control populations when aged on a HS diet for 3 weeks (Fig 4A-B). Clusters of co-regulated genes in each sex had similar expression patterns, consistent with what would be expected from coordinated regulation of signaling pathways. With the exception of the S4-C4 pairing, females exhibited a greater number of significant alterations in their transcriptome from their paired population. (S2 Table). As was the case for allele frequencies, the fewest differences in expression were seen between populations S2 and C2. Using principal component analysis (59), we clustered expression data from HS-fed flies and noted that there is modest clustering in expression profiles among the selected populations (S8 Fig). Among females, S3 and C3 had the most pronounced differences for the genes in PC1 (S8 Fig) and among males, S4 and C4 had the best separation over PC1, consistent with the large improvements in lifespan seen in the two Selected populations. These scores plots highlight the diversity among population pairs and suggest that the Selected populations exhibit similar expression trends, but limited direct overlap in specific DE transcripts, as was observed for our gene-level analyses (Fig 3A). Having identified an overabundance of SNPs in neuronal genes, we next assessed these pathways in greater detail.

**Fig 4.**
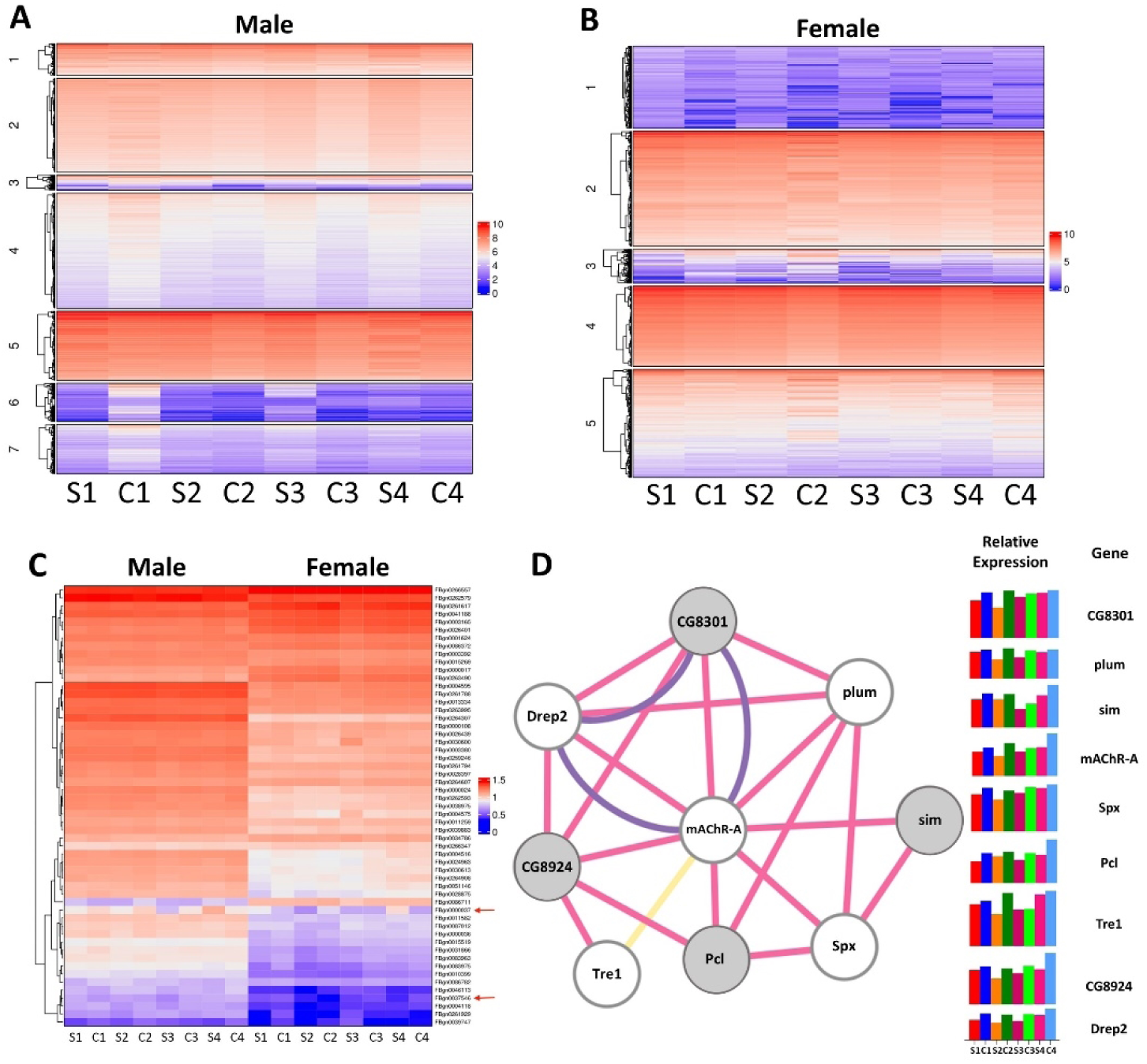
Differential expression profiles in HS-Selected flies suggest a role for the CNS. Flies were fed HS for 3 weeks and whole animal RNA was used for RNA-seq. Male and female data were analyzed separately due to the overwhelming influence of sex on gene expression (S7 Fig). (A-B) Expression is plotted as log-transformed median expression values. Heat maps after cluster analysis show differential expression patterns for all genes and genes with the same cluster typically have similar expression profiles. (A) shows males, and (B) shows females. (C) Heat map hierarchical clustering featuring genes in the ontology categories of neural development, learning and memory, and synaptic chemical transmission that were differentially expressed in a sugar- and population-specific manner. Males are on the left, females on the right. The arrows in (C) highlight the genes encoding the muscarinic acetylcholine receptors *mAChR-A* (FBgn0000037) and *mAChR-B* (FBgn0037546), which showed significant differential expression across selected populations. (D) Shows a gene network produced by Cytoscape showing differentially expressed genes (circles) connected to *mAChR-A*. Lines representing co-expression, purple co-localization, and yellow representing a shared protein domain. Genes in gray are those in the DNA binding gene ontology category. Bar graphs show log-transformed median expression values for males from each population.

### Neuronal function and cholinergic signaling candidate genes

Given the prevalence of reduced cognition upon aging and the allele frequency differences associated with cognition and neuronal function between the Selected and Control populations, we assessed the expression of genes in this category (Fig 4C). Several cholinergic signaling genes exhibited statistically significant allele frequency changes and differential expression between paired Selected and Control populations. Both nicotinic and muscarinic receptors were affected, and changes in these and acetylcholine regulators were observed across populations (S1 Table, S9 Fig). We thus assessed the expression of genes associated with neuronal function in the fly that were differentially expressed in at least one population. Here we observed patterns of differential expression in neuronal genes, with the muscarinic acetylcholine receptor *mAChR-A* standing out in particular (Fig 4C). This gene, which exhibited allele frequency differences in four comparisons, was down regulated in females in two of the Selected populations and in males in three of the Selected populations (S1 Table, Fig 4C-D). The *mAChR-A* gene encodes a G protein-coupled receptor (GPCR) orthologous to the human stimulatory muscarinic acetylcholine receptors M1, M3, and M5, which trigger intracellular calcium release upon ligand binding (60–62). mAChR-A is expressed strongly and specifically in the male and female adult brain (63). This gene was also found in PC1 for differential expression in both males and females (S8 Fig). Interestingly, several genes that exhibited similar expression patterns in our study have been previously described as having co-localization, co-expression, or shared protein domains with *mAChR-A* (Fig 4D). Several of these genes have been shown to be expressed in the brain and control neural physiology and behavior (37,64–69). These genes showed similar trends of expression across Selected and Control populations as *mAChR-A*, typically exhibiting lowered expression in the Selected populations (Fig 4D). Taken together, these data suggest that variation in *mAChr-A* and related genes contributes to adaptation to the HS diet and led us to measure behaviors that may have been beneficial to lifespan on a HS diet.

### *mAChR-A* alters lifespan and feeding behavior in a sugar-specific manner

To explore potential links between cholinergic signaling and longevity in relation to HS feeding, we selected *mAChR-A* because of its interesting differential expression pattern (Fig 4C-D) and changes in allele frequency in multiple populations (S1 Table). This receptor was targeted with RNA interference in the brain by crossing *syb-GAL4* with *UAS-mAChR-A*^RNAi^, hereafter referred to as “*mAChR-Ai”*. The control was a cross between the driver and the insertion site control genotype. Both the control and *mAChR-Ai* larvae were reared on a 5% dextrose-cornmeal-yeast-agar media and aged as adults on 0.15M or 1M sucrose. To address the role of the muscarinic acetylcholine receptor in longevity, we first measured lifespan. There was a highly significant 0.16 fold decrease in lifespan in *mAChR-Ai* males, but no significant difference was observed in transgenic females aged on the HS diet (Fig 5A,B) or for either *mAChR-Ai* males or females aged on 0.15M sucrose (Fig 5C,D), compared with the control genotype. This suggests that mAChR-A signaling is important for survival during HS feeding and operates in a diet-specific and sex-specific fashion.

**Fig 5.**
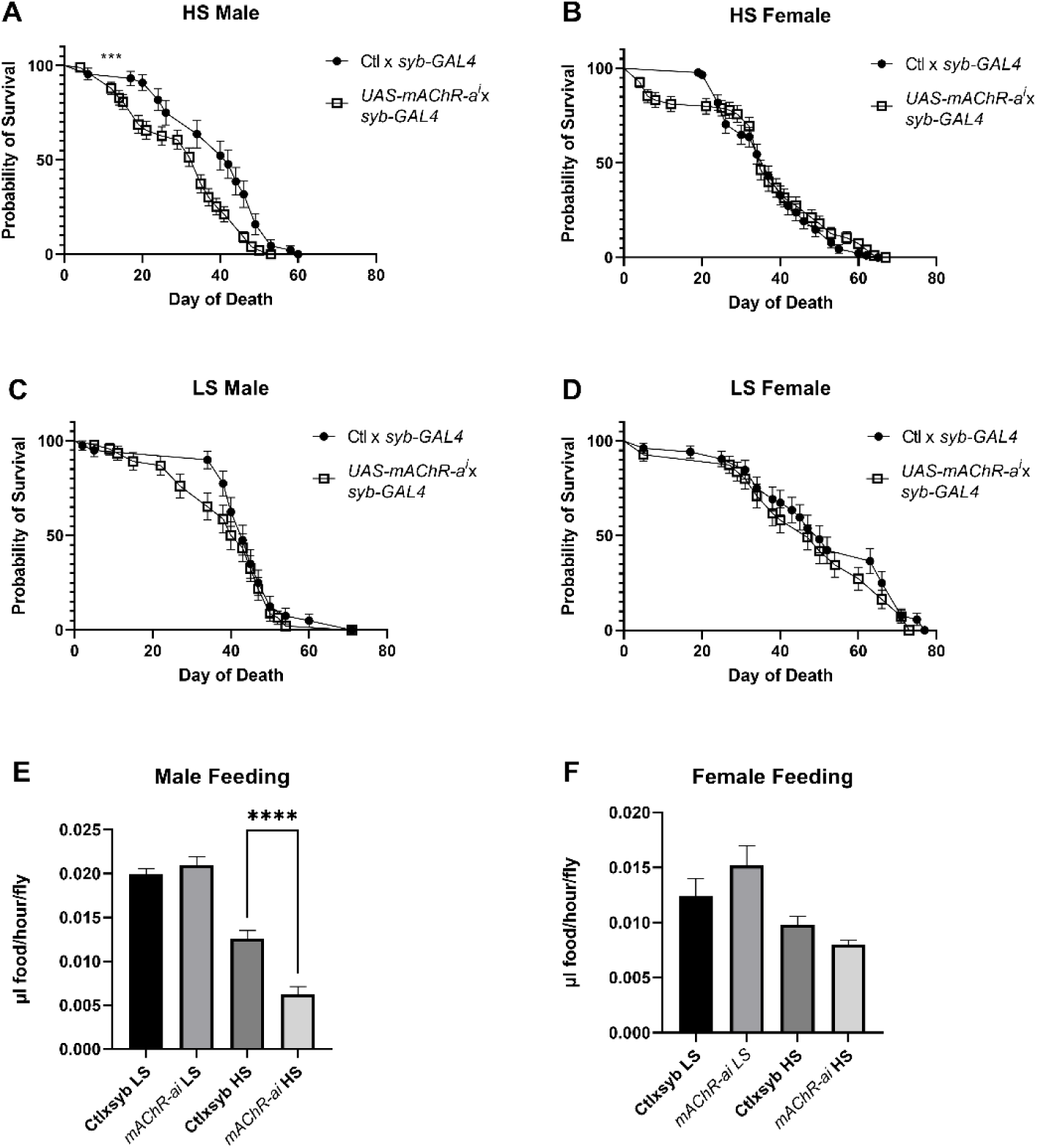
The type A muscarinic acetylcholine receptor promotes longevity and feeding in a sex- and sugar-specific fashion. (A, B) Neuron-specific RNAi targeting *mAChR-a* produced a significant reduction in survival of HS-fed males but not females. (C, D) There was no effect (P>0.19 for both) of this transgene on the median lifespan in control-fed flies. (E) Knockdown of *mAChR-a* reduced feeding in HS-fed but not control-fed males. (F) There was no significant effect of this transgene on feeding in females fed either diet. (A) *** P < 0.001 by log-rank Mantel-Cox test. (E) **** P < 0.0001 by one-way ANOVA.

Because the CNS controls feeding behavior and previous studies have shown HS affects feeding (64,65), which in turn can have profound effects on longevity, we measured the feeding rate of control and *mAChR-Ai* flies using a ‘Con-Ex’ assay (58). Control and *mAChR-Ai* animals were aged as adults on 0.15M or 1M sucrose for 3 weeks before being transferred to media supplemented with 2% FD&C Blue #1 dye for 2 h. We again observed diet- and sex-dependent effects, as *mAChR-Ai* significantly reduced feeding only in male flies aged on HS while there were no significant effects on consumption in control-reared males or in females on either diet (Fig 5E,F). This result might be expected given the higher expression of mAChR-A in males, compared with females, in both our whole-animal expression data and public adult brain expression data from Dr. Julian Dow and colleagues (S2 Table, (63)).

### Pharmacological inhibition of muscarinic signaling produces sugar-specific phenotypes

As the Selected population S3 exhibited alterations in *mAChR-A* and showed the most striking extensions in lifespan (a trend that continued into generation 15 and beyond), we attempted to validate the role of muscarinic cholinergic signaling in lifespan extension by treating this population and its paired control with atropine ((*RS*)-(8-Methyl-8-azabicyclo[3.2.1]oct-3-yl) 3-hydroxy-2-phenylpropanoate), a specific inhibitor of muscarinic receptors (70). To measure lifespan, mated adult flies were aged on HS food with or without atropine. Similar to *mAChR-Ai*, atropine produced sex-specific and diet-specific effects, and atropine also produced different results in the Control and Selected populations. The largest effect on lifespan observed in Control population C3 males, where it exhibited a 0.125-fold reduction in median day of death (P= 0.0009) on the HS diet (Fig 6A, atro) while the corresponding population S3 males were not significantly affected by atropine supplementation (P = 0.53, Fig 6A). Again, this tracked with expression, as the largest effect of atropine was observed in the flies with the highest expression of *mAChR-A,* population C3 males. The opposite effect was observed in the females where population S3 females exhibited a modest 0.06-fold increase in median day of death (P = 0.039, Fig 6B) while the Control C3 females were not significant affected (P= 0.09, no fold increase, Fig 6B). Because *mAChR-Ai* flies exhibited a feeding phenotype, we examined the effects of atropine on feeding rates (58). To measure the effects of atropine on feeding, mated adult flies were aged for 3 weeks on +/− atropine HS and LS food. Atropine did not have a significant effect on feeding for many of the comparisons on HS diets (Fig 6C-F), although a significant reduction in feeding was observed in Control population C3 females (Fig 6D).

**Figure 6.**
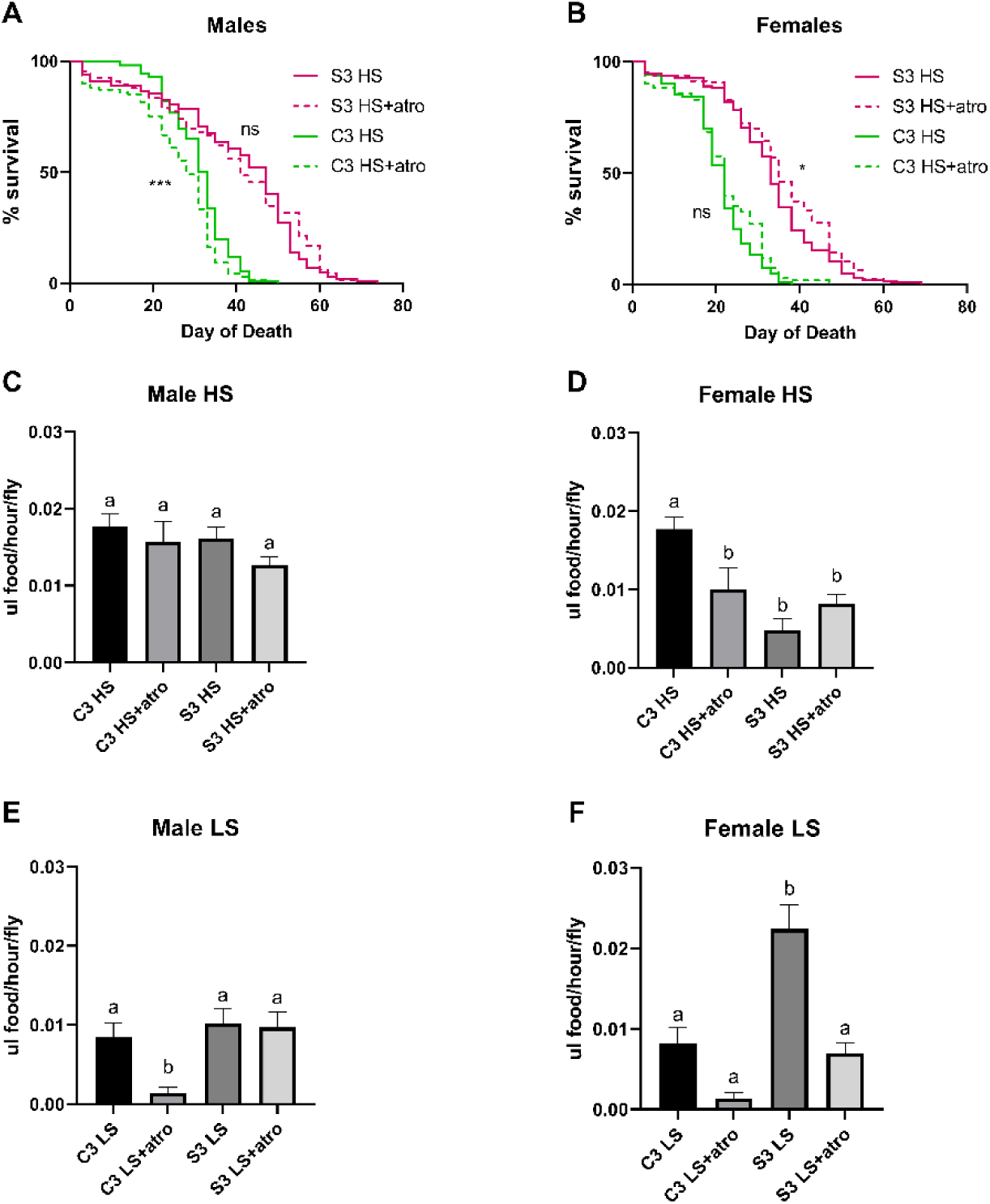
Supplementation with the muscarinic cholinergic antagonist atropine produces population, sex, and sugar-specific phenotypes. (A) Adding atropine (atro) to the HS diet reduced the lifespan of population C3 males, but not population S3 (P = 0.0009 and P = 0.53 respectively). (B) There was a slight improvement in female lifespan with atropine supplementation that reached significance for population S3 (P = 0.039) but not C3 (P = 0.0876). (C-F) Atropine reduced feeding in some genotype x diet combinations. (C) There was no effect of atropine on feeding in HS-fed males. (D) Atropine significantly reduced feeding in HS-fed C3 females (P = 0.0003). (E) On a control diet, C3 male feeding was inhibited by atropine (P = 0.005). (F) Atropine appeared to inhibit feeding in control-fed females of both populations; C3 0.15M female vs C3 0.15M atropine female P = 0.1, S3 0.15M female vs S3 0.15M atro female P < 0.0001. Significance is determined by one-way ANOVA and Tukey post hoc tests. Different letters indicate significant differences (P < 0.05).

Interestingly, atropine eliminated the dramatic difference in feeding between population S3 and C3 females on HS (Fig 6D). These sex-dependent results of atropine on feeding of the HS diet were consistent with differential expression of its target receptor, mAChR-A, in males from populations S3 and C3 (fed on the HS diet, compared to control-fed males, as shown on the heatmap in Fig 4C).

To further explore potential roles for muscarinic signaling in feeding, we tested atropine on the control (LS) diet. A more compelling trend toward reduced feeding was apparent in atropine-fed flies on the control diet (Fig 6E,F). Females from both populations and males from population C3 all reduced intake after atropine supplementation (Fig 6E,F). As for longevity, there was no effect of atropine on feeding in S3 males (Fig 6E). Females from population S3, who typically reduce feeding on the HS diet, no longer did so when atropine was present (Fig 6D,F P = 0.0003 via one-way ANOVA). Taken together, our data are consistent with a diet- and sex-dependent role for muscarinic acetylcholine receptor signaling in *Drosophila* feeding and longevity.

## Discussion

In this study, we used an evolve and resequence (E & R) approach to identify key genes and pathways in age- and overnutrition-associated pathophysiology phenotypes. Laboratory selection prompted a highly polygenic and sex-biased response across replicate populations. There was evidence for parallel evolution at the levels of biological processes, pathways, and gene families, although there was limited overlap in the SNPs and individual genes identified. Genes involved in cholinergic signaling, as well as a number of potentially interacting genes, exhibited significant differences in allele frequency and/or expression across selected populations. We verified one such gene with both genetic and pharmacological manipulations. A knockdown of *mAChR-A* reduced male lifespan and consumption behavior in a sugar-specific fashion. Furthermore, flies treated with atropine, an antagonist of muscarinic acetylcholine receptors, also showed sex-specific alterations in lifespan and feeding. These results suggest that our E & R approach was successful at identifying causal variants contributing to lifespan under the conditions used.

### Increases in lifespan

Evolve and resequence studies in Drosophila have been successful at producing rapid responses to selection and at mapping the genetic basis for a variety of complex traits (11,14,17–19,71–74). In the current study, we observed a robust phenotypic response to 10 generations of adaptation on two food media that correlated with polygenic changes in the genome and differential expression (Fig 2-4). By generation 10, all Selected populations exhibited significantly increased lifespan on both HS and LS diets, while Control populations had increased lifespan primarily on the LS (Fig 2, S1-4 Fig). Our results suggest that both Selected and Control populations underwent rapid selection for longevity, but the median lifespan extension was greater in magnitude in the Selected populations. The more dramatic response in the Selected populations is consistent with previous literature showing that stronger selective pressures elicit a more rapid response in the trait under selection (75–78). Strikingly, the males from Control populations C1 and C4 showed a modest yet significant increase in survival on HS food, despite being high-sugar naïve, after 10 generations (Fig 2). These data suggest that loci contributing generally to longevity may have undergone genomic changes in C1 and C4 that extend male lifespan on both diets, although the effect is attenuated on HS. Given these overlaps, a potential future avenue may be to explore loci linked with longevity across diverse environmental conditions.

### Evidence for parallel evolution

Understanding whether adaptation is predictable and can proceed via parallel evolution from standing variation is a major question in evolutionary genetics. Our results support the hypothesis that the predictability of evolution depends upon the hierarchical level being tested (79). All Selected populations showed significant increases in lifespan on HS as compared to Controls, suggesting parallelism at the level of overall fitness. Selected populations also showed a marked overlap in ontological categories based on significant changes in allele frequencies, providing support for parallel evolution at the levels of biological processes, organ systems, signaling cascades, and gene families. Perhaps most striking was the prevalence of CNS genes associated with neuronal development, behavior, and synaptic signaling. We observed a diverse array of acetylcholine receptors and related cholinergic genes with significant changes in allele frequencies in all of our Selected populations (S1 Table). All Selected populations had similar changes in the expression of genes functionally related to *mAChR-A* (Fig 4D).

Considering these parallels, it may be surprising to note that there was little evidence of parallel evolution at the polymorphism level. Only five genes overlapped across Selected populations (Fig 3A); although each gene did share multiple polymorphisms with significant changes in allele frequencies across all four population pairs (P < 1.0^-10^ for all Selected populations). Further, some, but not all, Selected populations exhibit an advantage over the Control on the LS diet (Fig 2, S1-4 Fig). This, coupled with the modest genetic overlap between Selected populations, suggests that the populations evolved unique solutions to our selective pressure at the individual gene and locus levels but many of the adaptive changes influenced similar biological processes. Given the rapid, high mortality rate in Selected populations at early generations, it is perhaps no surprise that the populations diverged at the genomic level. Beneficial alleles, especially for a highly polygenic trait like lifespan, might easily be lost due to chance under strong selection. These data are consistent with other studies showing that stochasticity plays an important role (79–81), especially in populations with extensive standing variation (79,82,83). Interestingly, among the neuronal GO terms, genes associated with development were the most enriched in our SNP gene lists despite our selective pressure being applied exclusively to adults. Taken together, these data suggested to us that the brain might have been particularly responsive to our selective pressure, whereas shifts in metabolic flux may have complex or antagonistic effects that make it difficult to achieve overall benefits to development, lifespan, and healthspan.

### mAChR

One of the genes associated with learning, memory, and synaptic transmission, *mAChR-A,* provided strong evidence for parallel evolution as it showed significant differences in allele frequencies in multiple Selected populations as well as reduced expression in Selected males from population S3, compared with Control population C3 (Fig 3A, Fig 4C, S1 Table). A network analysis of one cluster of differentially expressed genes in our male dataset revealed genes connected to *mAChR-A* through previously established co-expression or co-localization. Like *mAChR-A*, expression of these genes decreased in males across all of our Selected populations. Both *polycomb-like (Pcl*) and *single-minded* (*sim*) have been shown to be involved in behavior likely to be relevant to lifespan and healthspan during HS overnutrition. The gene *sim* has been associated with the development of the midline cells of the embryonic nerve cord and the brain central complex (67–69) and *sim* mutants exhibit defective locomotion behavior (66). Moreover, it has been demonstrated that *Pcl*, in its role as part of the Polycomb Repressive Complex 2.1, is an integral epigenetic regulator of sugar-dependent taste and feeding in *D. melanogaster* (37,64,65). This complex binds the chromatin of sweet-sensing gustatory neurons, repressing transcription involved in sweet taste; this thus modulates appetitive behavior and promotes obesity (37,64,65). Reduced *Pcl* expression might therefore benefit Selected flies by attenuating feeding.

The *Drosophila* muscarinic acetylcholine receptors or mAChRs comprise a family of three G-protein coupled receptors (type A, B, and C) - two of which, A, and B, are strongly expressed in the central nervous system (63). Mammals possess two classes of mAChRs, M1-type and M2-type, with *mAChR-A* sharing the closest homology with the excitatory M1-types (60–62). Recent studies have shown both stimulatory and inhibitory roles for muscarinic signaling in different types of *Drosophila* neurons (84,85). In addition, *mAChR-A* has shown age-dependent, spatially restricted expression patterns in neurons of the adult *Drosophila* brain (86) (86). These data, coupled with a general decrease in expression of mammalian muscarinic acetylcholine receptors with age suggest that *mAChR-A* may play a conserved role in senescence (87–94). In the present study, *mAChR-Ai* produced a reduction in both lifespan and consumption in HS-fed males (Fig 5). Because our populations exhibited reduced *mAChR-A* expression in response to selection correlating to their increased lifespan, we anticipated that knockdown of *mAChR-A* would improve fitness on HS. The reduction in lifespan we observed in the knockdown flies may result from a critical effect on feeding or other cholinergic-regulated behaviors that negatively affect lifespan. It may be too simplistic too expect basic whole-gene manipulation to recapitulate a complex role. Selected flies may have co-adapted protective genes that help to maintain fitness in the face of reduced feeding, a genetic context that is impossible to replicate in a transgenic animal. To complement the knockdown, we administered the S3-C3 population pair atropine, an inhibitor of mAChR (Fig 6). Here, we observed a trend toward marginal improvements in female lifespan, although the C3 population males (with higher expression of *mAChR-A*) exhibited a significant reduction in lifespan, similar to the knockdown flies. Given that there is a complex interaction between feeding and nutritional geometry that may cause variable intake of the drug, water, and nutrients, it is difficult to interpret these paradoxical results to determine exactly how atropine impacts lifespan. In addition, mAChRs are broadly expressed at different types of synapses and may regulate developmental and behavioral processes that these data are unable to capture (60,85,88,90–92,95–97). Furthermore, atropine targets all mAChRs and thus may have affected the type-B and type-C receptors in a way the knockdown did not.

Curiously, the *mAChR-Ai* knockdown exhibited a sex-specific phenotype in that only males showed a reduction in lifespan and feeding. This phenotype is consistent with *mAChR-A* having higher expression in *Drosophila* males in our study (Fig 4C). The creators of FlyAtlas2, Leader et al., observed higher expression of *mAChR-A* and *mAChR-B* in adult male, compared to female, brains (63). Xia et al. further demonstrated that the type-C mAChR is more highly expressed in male heads than in female heads (90). *Drosophila* males may therefore be more sensitive to the interaction between mAChR signaling and high sugar due to their higher endogenous expression of these receptors. It has also been reported that mammalian mAChRs display sex-specific expression patterns, and drugs targeting the receptors differ in their effects by sex (91–93). Taken together, these data are consistent with other models where the effects of genotype on aging are sex- and brain-dependent. These results were also consistent with a broader trend of sex-specific quantitative traits present in the Selected and Control populations.

### Sex-specific effects of adaptation

The Selected populations exhibited sex-specific alterations at both the expression (Fig 4) and phenotypic (Fig 2) hierarchical levels. Males showed a greater increase in longevity than females after adaptation, compared to both Gen 0 and paired Controls. This ran counter to our expectations, given the dietary nature of the selective pressure, as mated female *Drosophila* have previously been shown to consume more than males (58,98). Females died more quickly on the HS diet, so that stronger selective pressure was applied to females, compared with males (Fig 2). Therefore, one might assume that the HS diet would have the most detrimental effects on females and elicit a stronger adaptive response. By contrast, our findings, which were consistent across genotypes, suggest males exhibit a greater response to HS than do females (Figs 2, 5, 6). This is consistent with previous studies showing that male flies fare worse during overnutrition (39,40,99,100).

Sex differences in the response to selection may also be a result of our unisex aging approach; cohousing flies reduces their lifespan (101). Both males and females suffer fitness costs during the courtship process (101–105). Courtship behavior and the production of ejaculate are energetically costly for males. Females, likewise, suffer significant energy costs during courtship and during oocyte and yolk production (102–105). Thus, unisex housing post-mating may have imposed a unique type of sex-specific selection pressure. In the absence of females, males will not engage in the cost of courtship. Females, by contrast, will continue to produce eggs even in the absence of males (though this declines with time). Therefore, we expected the influence of mating on lifespan to be relatively low in our study. In addition to the demands of courtship, mating, and egg/sperm production, males and females experience interlocus sexual conflict (101–103,105). Here, male- or female-biased selection may result in loci that confer fitness advantages for one sex at a cost to the other sex (102,103,105–108). For example, male *Drosophila* courtship behavior involves persistent “chasing” and aggressive attempts at mounting that are detrimental to female survival (101,104,105). Such behaviors in males are energetically costly to maintain, and the alleviation of such costs, as well as other types of conflict, may have contributed to the sex biases we observed in survival. The underlying mechanisms behind these sex differences will require future studies to elucidate them.

## Conclusions

In summary, our data indicate a robust phenotypic and genotypic response to selection for long-lived flies in two dietary environments using *Drosophila.* All replicate Selected populations exhibited increased lifespan on HS accompanied by genomic changes at many loci. Interestingly, we observed parallel adaptation at the pathway level across replicates with limited overlap at the gene or nucleotide levels. We identified a number of neuronal genes with putative roles in the physiology of aging on HS. Once fully characterized, *mAChR-A* and associated genes may provide important insight into the physiological mechanisms by which organisms maintain fitness over the lifespan in the face of overnutrition.

## Acknowledgements

The funding for this research came from Binghamton University and National Institute of General Medical Sciences of the National Institutes of Health (1R15GM128158-01). We would like to thank Julien Ayroles of Princeton University for providing the starter population, the Bloomington Drosophila Stock Center for fly stocks, and the McDonnell Genome Institute at Washington University in St. Louis for their sequencing services. We would also like to thank Matthew Pereira, Christie Santoro, and Utsav Nyachhyon for their help maintaining populations.

## Supplementary Figures

**S1 Fig.**
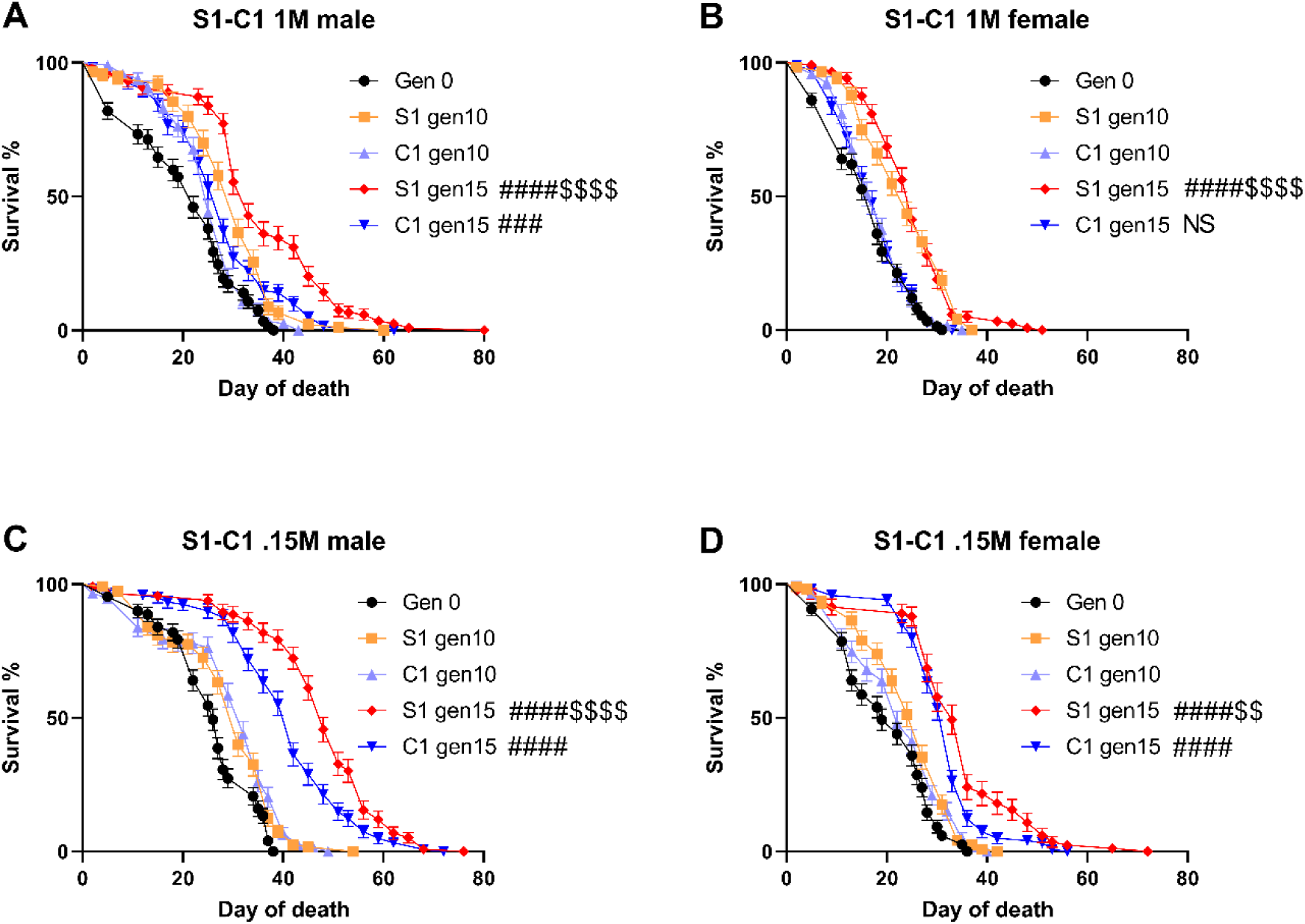
Population survival of S1-C1 pair with confidence intervals. In cohorts of 30, 120 recently eclosed mated males (A, C) from or females (B, D) were aged on 1M or 0.15M sucrose diets. Kaplan-Meier estimator curves for Generation 0 in black, Control populations are shown in cool colors, Selected populations in warm colors. The significance corresponding to number of symbols is as follows: *P < 0.05, **P < 0.01, ***P < 0.001, ****P < 0.0001 by log-rank Mantel-Cox. # represents the significance for generation 15 vs generation 0. $ represents the significance for the Selected population vs Control population at generation 15.

**S2 Fig.**
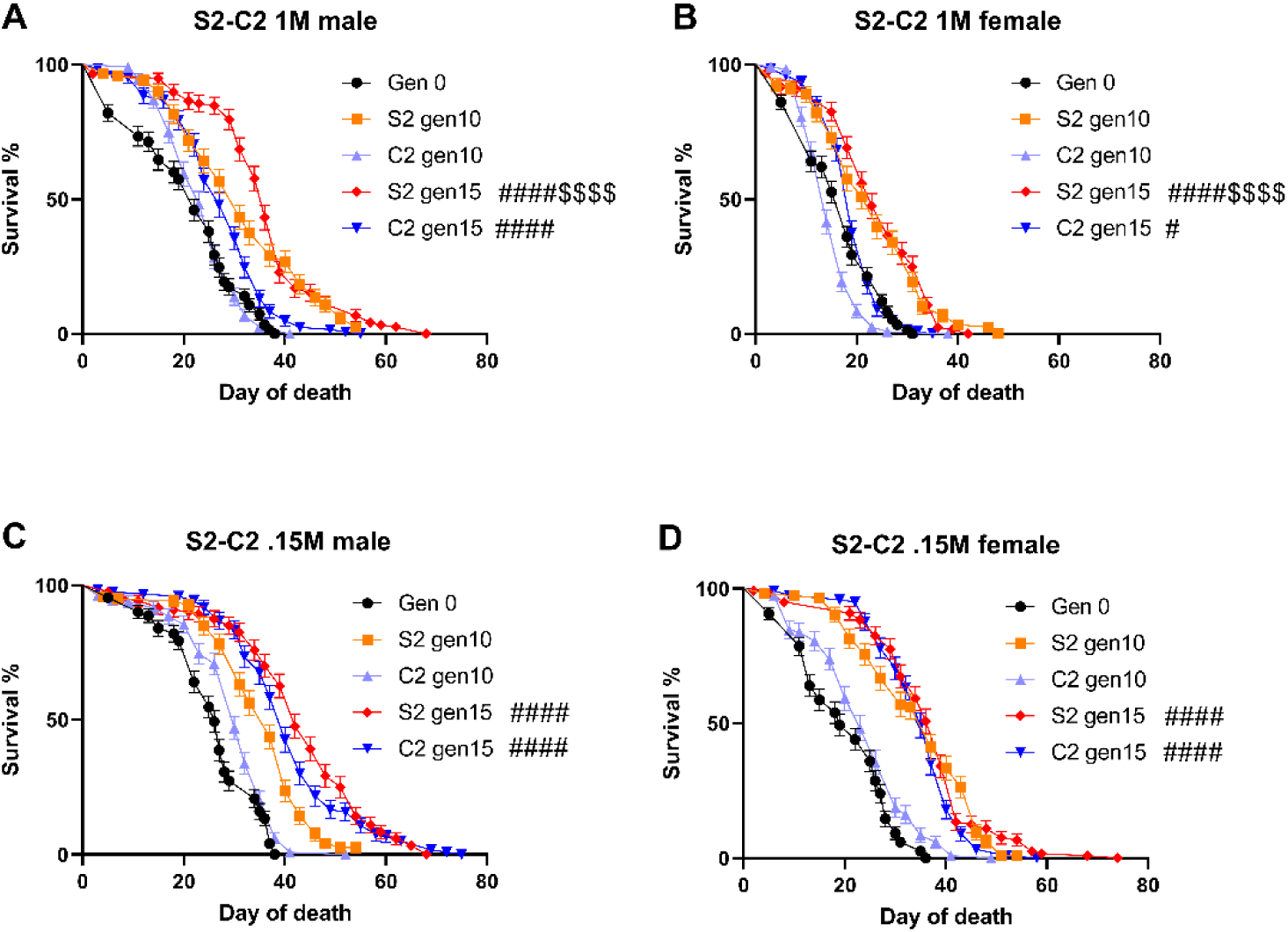
Population survival of S2-C2 pair with confidence intervals. In cohorts of 30, 120 recently eclosed mated males (A, C) from or females (B, D) were aged on 1M or 0.15M sucrose diets. Kaplan-Meier estimator curves for Generation 0 in black, Control populations are shown in cool colors, Selected populations in warm colors. The significance corresponding to number of symbols is as follows: *P < 0.05, **P < 0.01, ***P < 0.001, ****P < 0.0001 by log-rank Mantel-Cox. # represents the significance for generation 15 vs generation 0. $ represents the significance for the Selected population vs Control population at generation 15.

**S3 Fig.**
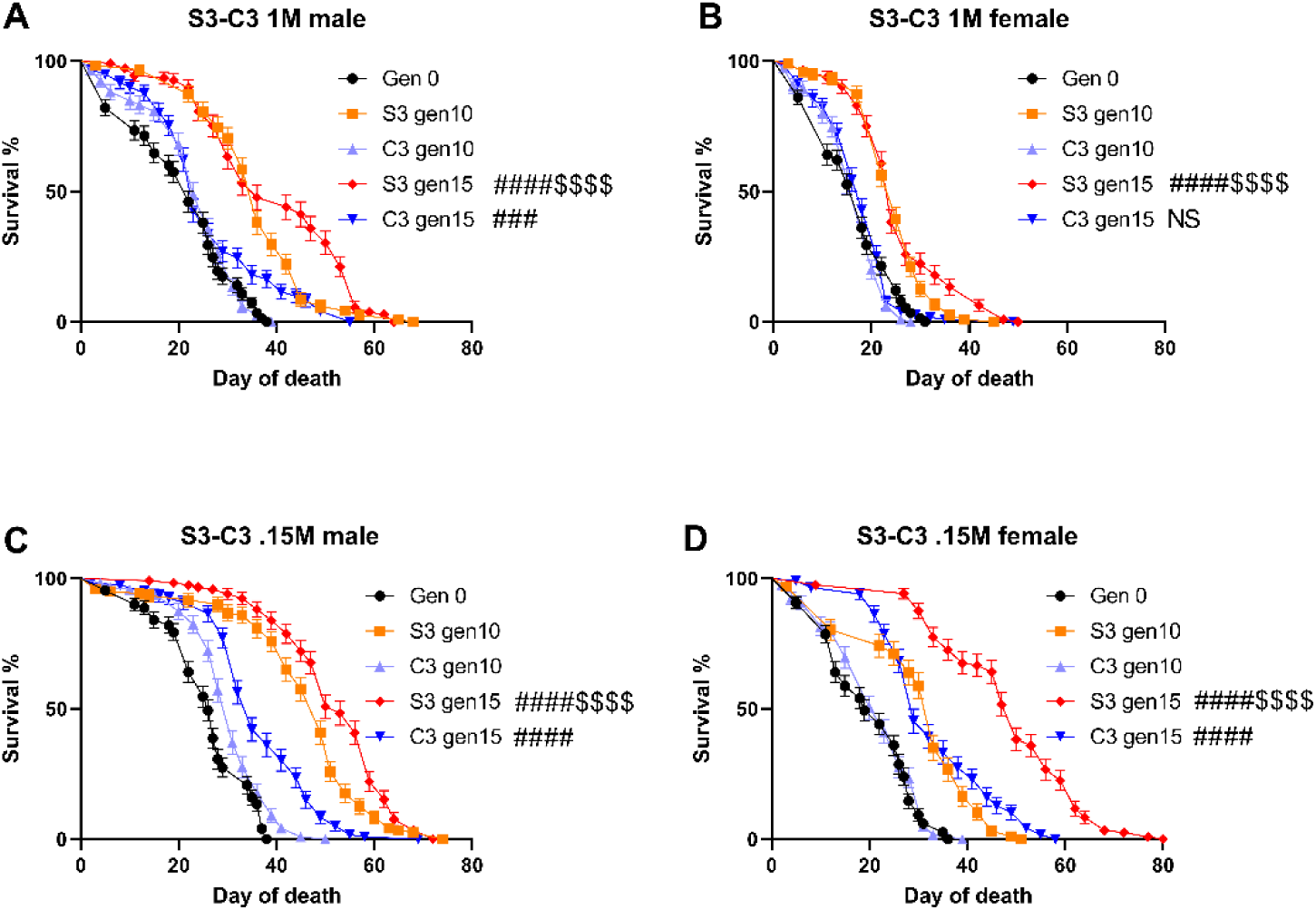
Population survival of S3-C3 pair with confidence intervals. In cohorts of 30, 120 recently eclosed mated males (A, C) from or females (B, D) were aged on 1M or 0.15M sucrose diets. Kaplan-Meier estimator curves for Generation 0 in black, Control populations are shown in cool colors, Selected populations in warm colors. The significance corresponding to number of symbols is as follows: *P < 0.05, **P < 0.01, ***P < 0.001, ****P < 0.0001 by log-rank Mantel-Cox. # represents the significance for generation 15 vs generation 0. $ represents the significance for the Selected population vs Control population at generation 15.

**S4 Fig.**
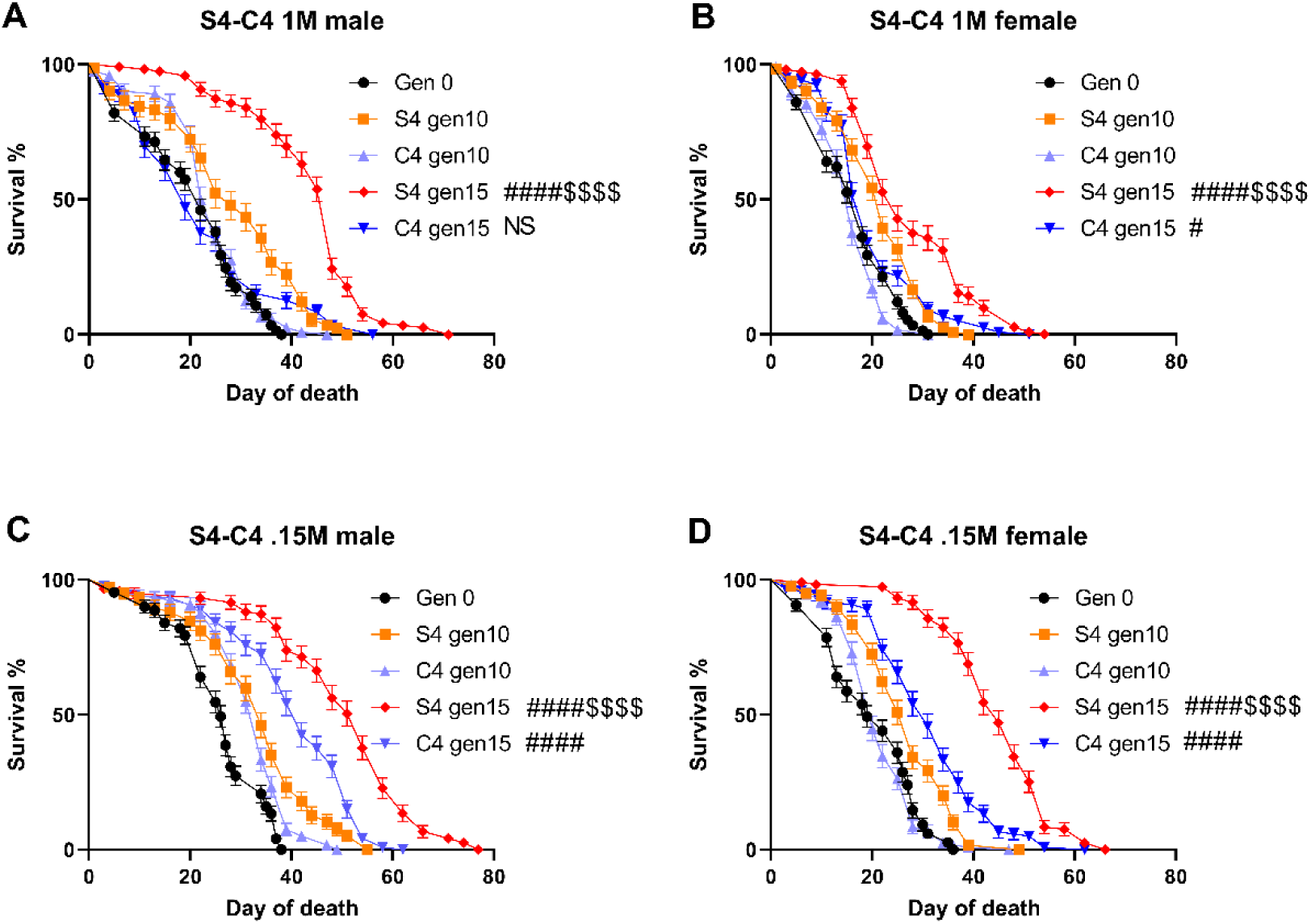
Population survival of S4-C4 pair with confidence intervals. In cohorts of 30, 120 recently eclosed mated males (A, C) from or females (B, D) were aged on 1M or 0.15M sucrose diets. Kaplan-Meier estimator curves for Generation 0 in black, Control populations are shown in cool colors, Selected populations in warm colors. The significance corresponding to number of symbols is as follows: *P < 0.05, **P < 0.01, ***P < 0.001, ****P < 0.0001 by log-rank Mantel-Cox. # represents the significance for generation 15 vs generation 0. $ represents the significance for the Selected population vs Control population at generation 15.

**S5 Fig.**
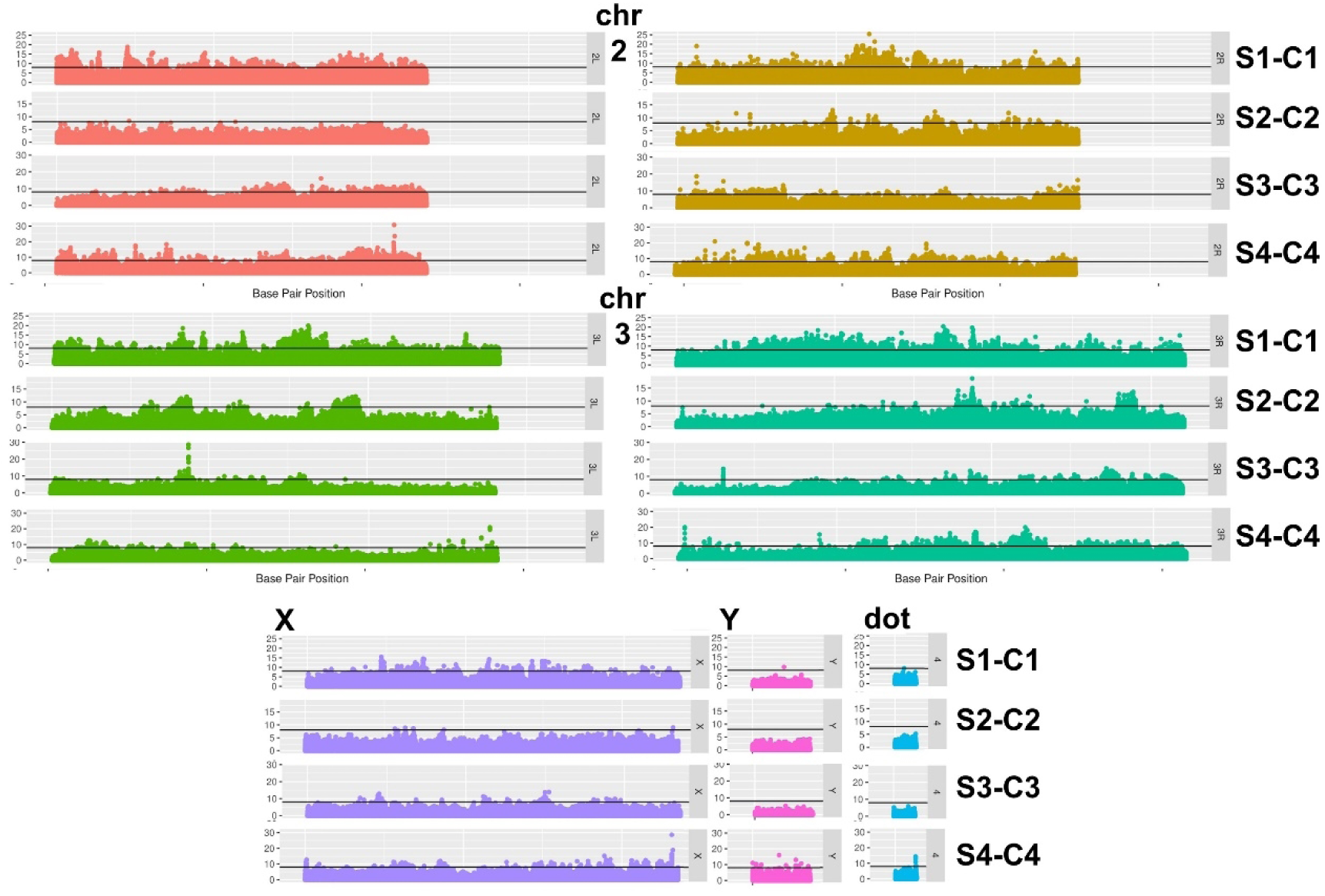
Pooled whole-genome sequencing shows changes in allele frequencies after ten generations of selection. Manhattan plots show SNPs identified as significant deviations in allele frequencies between Selected populations and Control populations via a Fisher’s exact test. Chromosomes are stacked for each of the different replicated comparisons. The horizontal line for each chromosome represents the P value cutoff (P < ∼10^-8^).

**S6 Fig.**
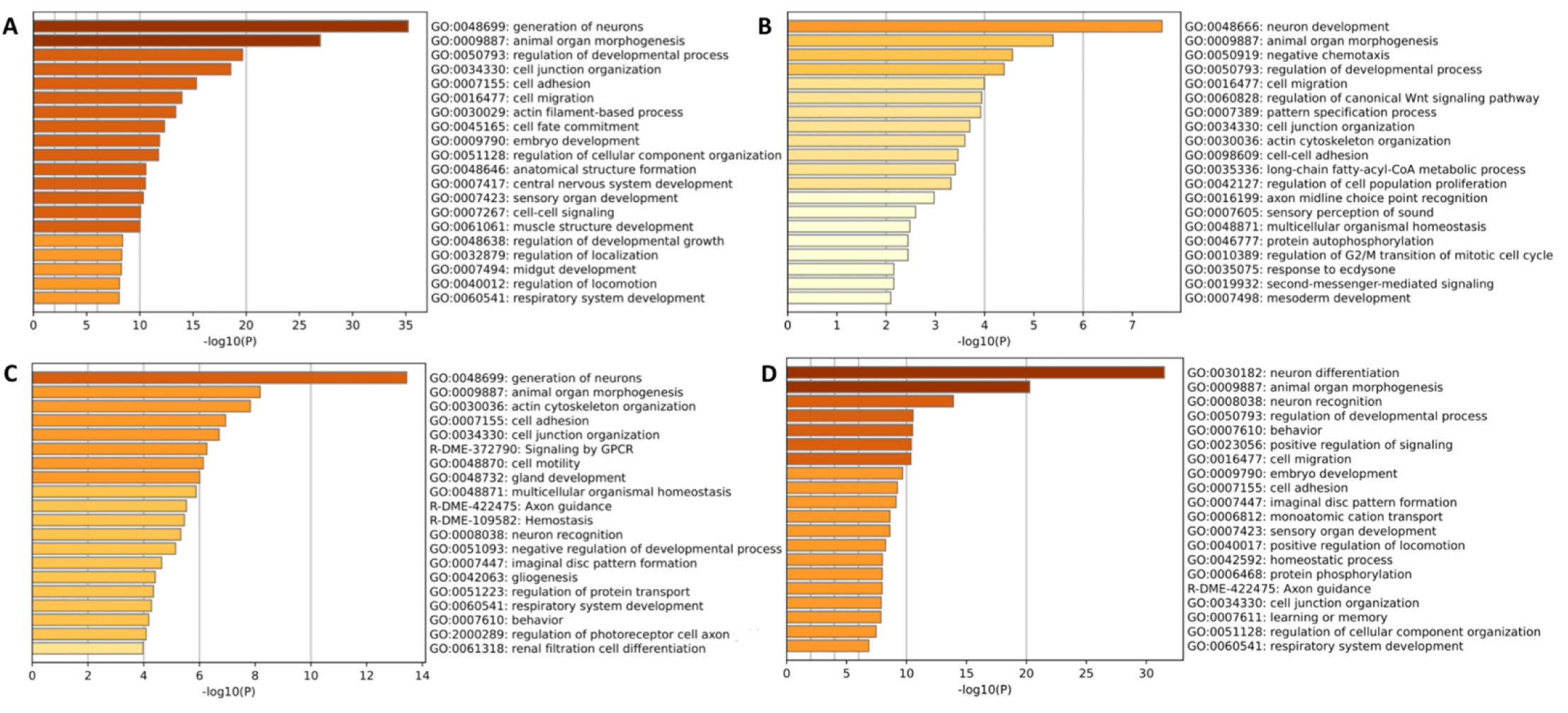
Gene ontology enrichment analysis of population pairs. Figures show top 20 enriched ontological categories obtained from significantly differentiated loci (by Fisher’s exact test) in (A) Populations S1-C1, (B) Populations S2-C2, (C) Populations S3-C3, and (D) S4-C4 population pairs. Terms with P < 0.01 and enrichment score >1.5 were counted as significant categories (ref for Metascape: https://pubmed.ncbi.nlm.nih.gov/30944313/).

**S7 Fig.**
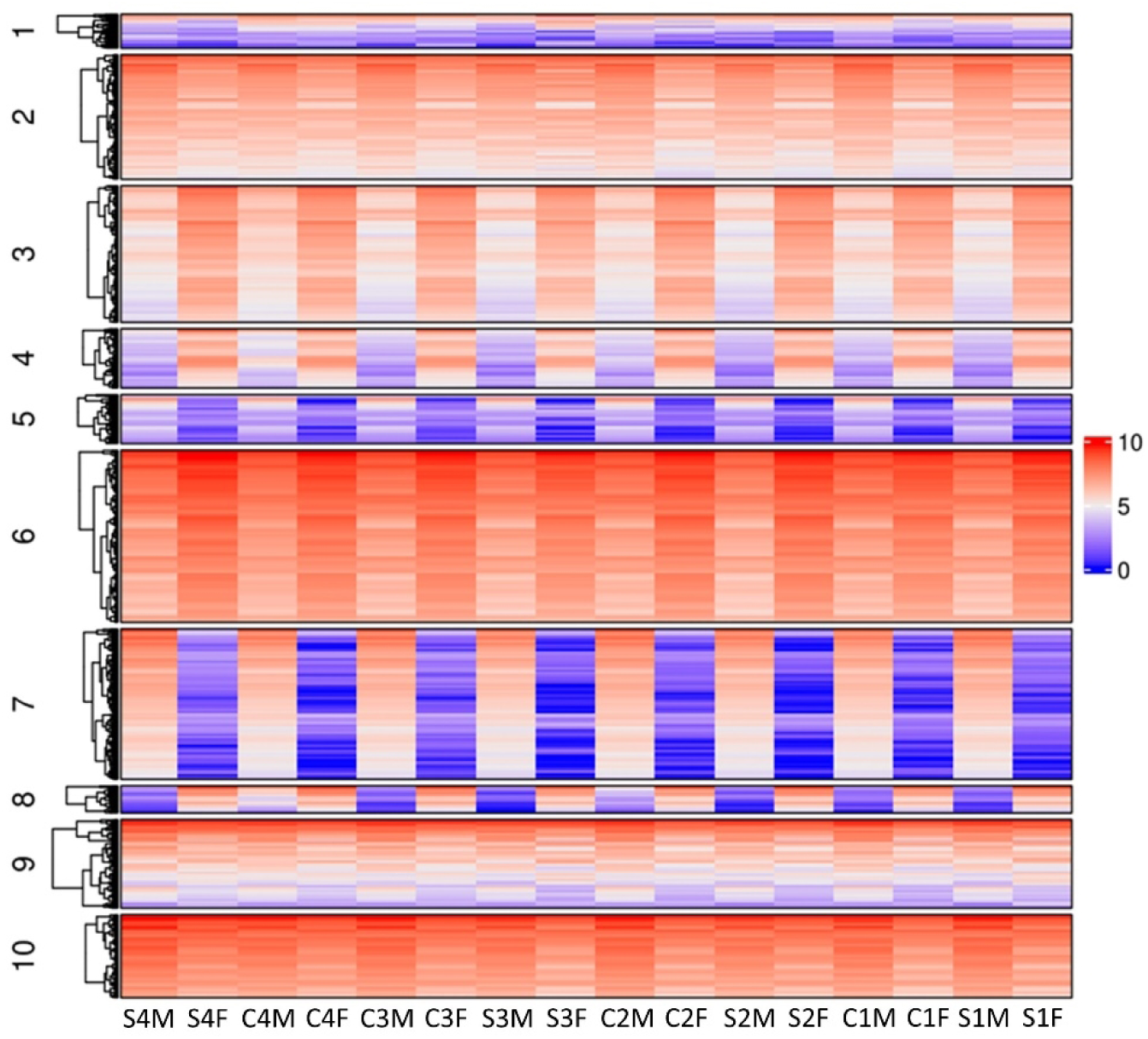
Differential expression profiles in HS-selected flies. Flies were fed HS for 1 week and whole animal RNA was used for RNA-seq. Cluster analysis of the median expression values from all populations for both males and females was conducted using a mixture of multivariate Poisson log-normal distributions. The models with number of clusters ranging from 1 to 15 were fitted and the model-selection criteria selected a model with 10 clusters. Log-transformed median expression profiles of all genes in all ten clusters are visualized using a heatmap.

**S1 Table:**
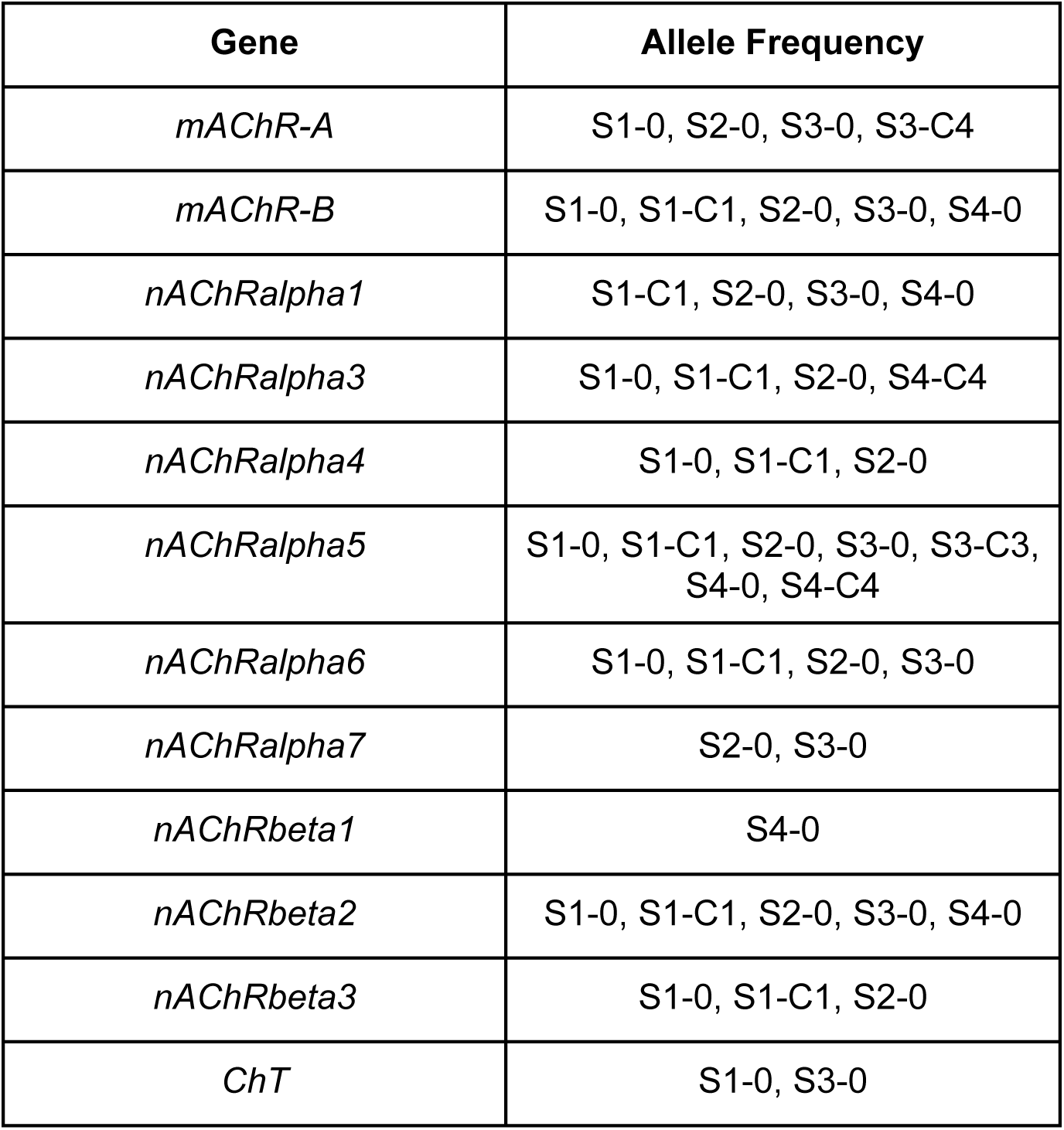
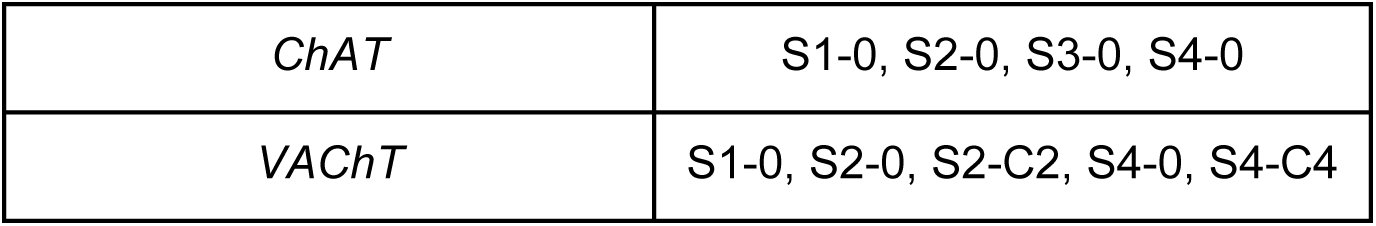
Cholinergic genes with significant changes in HS-selected fly populations. A diverse array of genes involved in cholinergic signaling (pre- and postsynaptic) were identified by both changes in expression and allele frequency in response to selection. Populations listed under allele frequency possess significant changes in allele frequency from either generation 0 (X-0) or from their paired control population at generation 10 (X-Y). ChT, choline transporter; ChAT, choline acetyltransferase; VAChT, vesicular acetylcholine transporter.

**S8 Fig.**
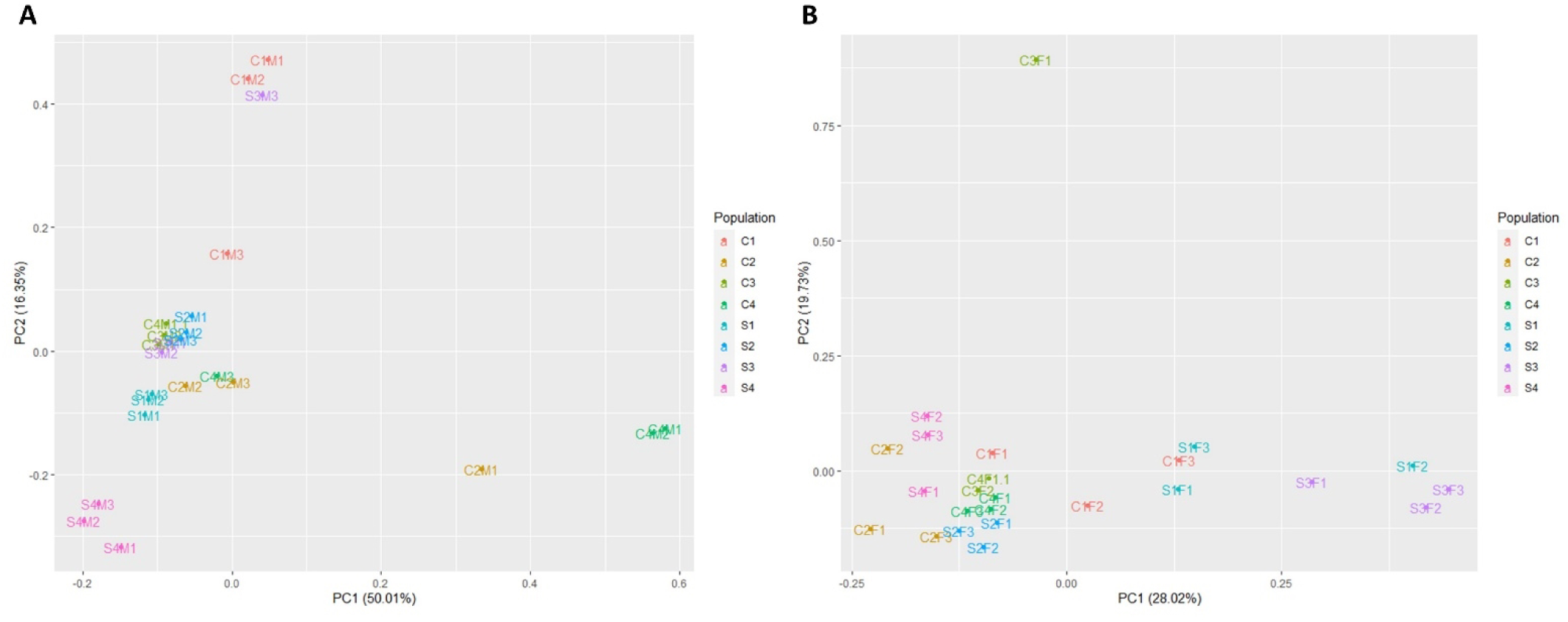
Populations differentiate by gene expression at generation 10. Adult male (A) and female (B) flies were fed HS for 1 week and whole animal RNA was used for RNA-seq. Principal component analysis (PCA) scores plots show the variance between normalized counts per million (CPM) of 3 replicates from control and selected populations. Colors represent populations.

**S9 Fig.**
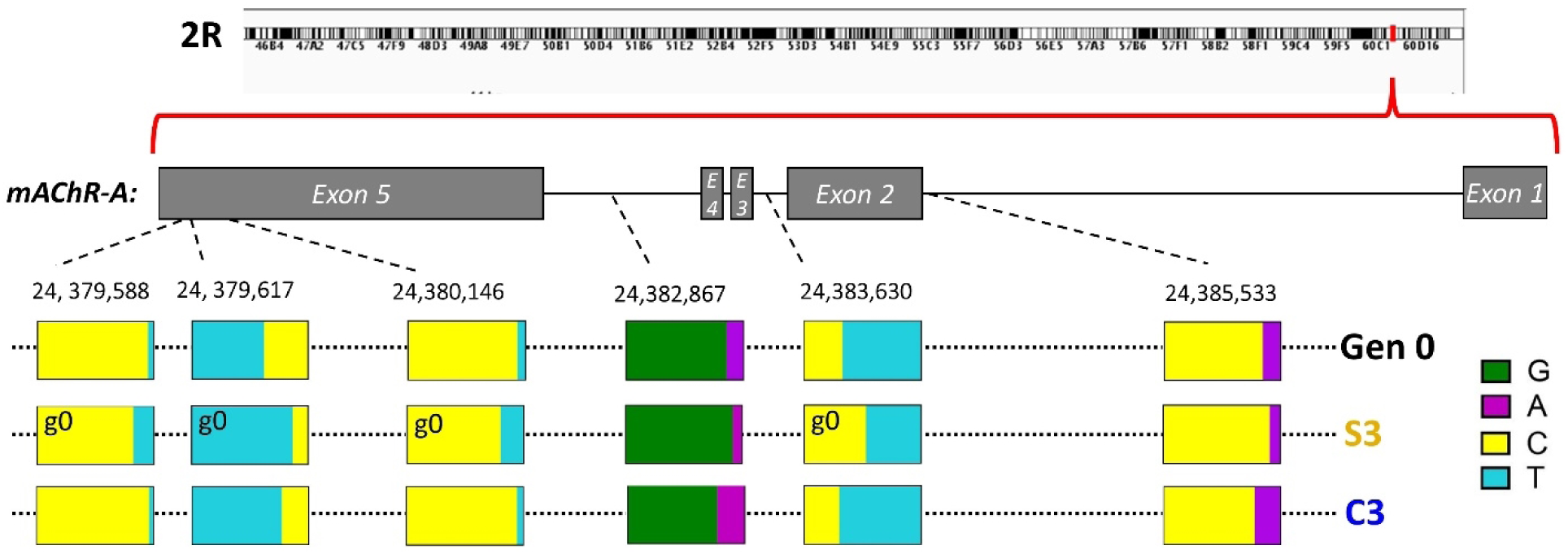
Changes in allele frequency in *mAChR-A* observed in populations S3 and C3. The top six changes ranked by lowest P value using a Fisher’s exact test all differed with P < 10^-5^ between S3 and C3. g0 denotes differences that reached the same degree of significance from generation 0. Generation 0, Selected population S3, and its corresponding Control population C3 are stacked to illustrate the major and minor allele frequencies.

